# Intrauterine Growth-Restricted Pregnant Rats, from Hypertensive Placental Ischemic Dams Display Preeclamptic-like Symptoms: A New Rodent Model of Preeclampsia

**DOI:** 10.1101/2024.02.07.579407

**Authors:** Jonna Smith, Madison Powell, Whitney Cromartie, Savanna Smith, Kylie Jones, Angie Castillo, Jordan Shaw, Joseph Editone, Ahfiya Howard-Cunningham, Robert Tatum, Alex Smith, Brandon Fisher, George W. Booz, Mark Cunningham

## Abstract

Preeclampsia (PE) is characterized by de novo hypertension (HTN) and is one of the primary causes of intrauterine growth restriction (IUGR). PE is associated with placental ischemia, decreased nitric oxide (NO) bioavailability, oxidative stress (OS), and organ damage in the kidneys and brain. This study aims to characterize a new model of PE using IUGR rats from hypertensive placental ischemic dams. It is hypothesized that pregnant IUGR rats from hypertensive placental ischemic dams will have elevated blood pressure (BP), OS, and organ damage.

**Methods:** Pregnant Sprague Dawley rats are divided into 2 groups: normal pregnant (NP) and reduced uterine perfusion pressure (RUPP) hypertensive placental ischemic dams. Offspring from NP and RUPP dams were mated at 10 weeks of age to generate pregnant IUGR (IUGR Preg; n=3-8) and pregnant CON (CON Preg; n=3-6) rats. BP and other markers of PE were evaluated during late gestation.

**Results:** Pregnant IUGR rats had elevated BP and systemic OS, as demonstrated by higher trending 8-isoprostanes and lower circulating antioxidant capacity. Maternal body weight of pregnant IUGR rats and their pups’ weights were decreased, while the brains were enlarged. Brain OS was elevated, with a rise in hydrogen peroxide (H_2_O_2_) and heat shock protein 1 (HSP- 1), along with lower Manganese Superoxide Dismutase (MnSOD) and antioxidant capacity.

**Conclusion:** Pregnant IUGR rats, born from hypertensive placental ischemic dams, have HTN and increased systemic and brain OS, with larger brain sizes and smaller pups. Pregnant IUGR rats exhibit an preeclamptic-like phenotype, which suggests a new epigenetic model of PE.

## INTRODUCTION

Preeclampsia (PE) a pregnancy complication syndrome that is generally defined by new-onset hypertension (HTN) usually occurring after 20 weeks of gestation [1]. In the clinic PE is diagnosed when there is hypertension and organ damage and dysfunction. Organ damage and dysfunction in PE patients is commonly presented as proteinuria, thrombocytopenia, increased liver enzymes, severe headaches, cerebrovascular dysfunction, and changes in vision [2–4]. Furthermore, Daughters of women who had PE during pregnancy are twice as likely to develop PE compared with women from normal pregnancies [5, 6].

PE is a syndrome with a wide variety of manifestations, timing, and severity [7]. PE causes end-organ damage, especially in the kidneys and brain, of both the mother and fetus [1]. PE is a major cause of maternal and fetal morbidity, mortality, intrauterine growth restriction (IUGR), and pre-term birth [4, 8–10]. PE affects ∼5-10% of all births in the USA each year [1].

IUGR can occur when there is a reduction in placental blood flow, causing the restriction of oxygen and nutrients to the fetus. This usually results in fetal growth retardation and the delivery of small-for-gestational-age babies [11]. PE affects both the mom and the fetus before, during, and after birth. Furthermore, PE increases the risk of cardiovascular disease, cerebrovascular disease, and renal disease for both mom and child later in life [1, 8, 12].

The origins and etiology of PE are not known, and there is no cure for the disease, except for delivery of the fetoplacental unit [4]. The pathophysiology of PE, in both clinical and animal studies, is associated with placental ischemia [4, 11, 13–15], decreased nitric oxide (NO) bioavailability, increased OS, and anti-angiogenic factors (such as soluble FMS-like tyrosine-1), along with genetic predisposition [4, 11, 16, 17]. To examine the pathology of PE and to derive new therapies against the development of PE, several rodent models have been created through pharmacological, surgical, and/or genetically induced alterations[11, 16, 18–21]. These models have pushed the field forward and provided significant insight into the pathophysiology of PE. However, these models are limited, because rodents, like many other animal species, do not naturally develop PE and require outside interventions to generate PE. Although these models provide a wealth of knowledge about PE, none of these models occur without intervention.

Few studies have examined pregnant rodents from 2^nd^ generation PE pregnancies, but none have done an extensive study, despite their increased risk of PE during pregnancy [22–24]. Therefore, in this study, we wanted to explore the pregnancy phenotype of rat offspring born from PE dams. We hypothesize that pregnant IUGR rats born from hypertensive placental ischemic dams will have HTN, reduced NO bioavailability, and elevated OS. The objective of this study is to characterize this new model of PE with adult pregnant IUGR rats from hypertensive placental ischemic dams.

## METHODS

### Animals

Timed-pregnant Sprague Dawley rats were purchased from Envigo (Indianapolis, IL) to produce offspring (both control and IUGR), which were mated and evaluated in this study. The animal protocols and handling methods were approved by Institutional Animal Care and Use Committee (IACUC) at the University of Mississippi Medical Center, #1543, where we conducted our animal experiments. The rats were housed in temperature-regulated rooms with 12-hour light-dark cycles. Food and water were administered ad libitum to all rats. All pregnant rats were allowed to give birth naturally and were weaned for three weeks. Animal experiments were conducted under the guidelines from the National Institute of Health (NIH) for the use and care of animals. The Western blots and ELISAs were performed at the University of North Texas Health Science Center.

### Pregnant IUGR Rats from Placental Ischemic Hypertensive Dams

All IUGR rats were derived from the placental ischemic hypertensive reduced uterine perfusion pressure (RUPP) model of PE; and from different RUPP dams. Note, that RUPP model is a commonly used surgical model of PE to generate IUGR offspring, which experience placental ischemia and are born with low birth weights [8, 11, 25]. Our P1 preeclamptic pregnant dams were separated into two groups: normal pregnant (NP) and RUPP groups. On gestational day 14, the RUPP dams received the RUPP surgery, as described by us and others [4, 11, 13, 14, 21, 25–27]. No surgical/sham procedures were performed on the NP dams, as previous studies from our labs and others showed no differences in sham vs NP control rats [28, 29]. Both groups were allowed to progress through their pregnancy and deliver naturally.

IUGR and CON rats were then separated by sex and group. At 10-12 weeks of age, the F1 offspring were mated to create two groups: Pregnant IUGR (IUGR Preg; n=3-8) and control pregnant (CON Preg; n=3-6) rats. Pregnant IUGR rats were produced by mating IUGR females to CON males and IUGR females to IUGR males. Whereas the pregnant CON rats were produced by mating CON females to CON males and CON females to IUGR males. Note, all offspring (IUGR and CON), used generate our pregnant rats, were from different dams. BP, body, organ (placenta, heart, kidney, and brain), and pup weights were recorded on gestational day 19 (late pregnancy, where most of the pathology of PE occurs in patients). Tissue and plasma samples were stored at -80°C to evaluate systemic and local levels of NO bioavailability and OS, using colorimetric biochemical assays and Western blots.

### Mean Arterial Pressure (MAP), Plasma, and Organ Collection

MAP measurements were performed on gestational day 19, the day after carotid catheterization, as described by us previously [25, 30–35].

### Plasma, Organ Collection, & Storage

Pregnant CON and IUGR rats were euthanized on gestational day 19, in which the blood and organs (kidney and brain) were weighed, collected, and stored at -80°C. Organs were later homogenized in 1:10 lysis buffer for Western blots or a 1X PBS or assay-specific buffer (50 mg: 500 μL buffer) for assay usage. The homogenate was then centrifuged at 10,000 g for 15 minutes to obtain the supernatants. Protein amounts of all supernatants were determined by bicinchoninic acid (BCA) protein assays and used to normalize all experimental values.

### Nitric Oxide (NO) Bioavailability

NO bioavailability was evaluated in the plasma, kidneys, and brains of pregnant CON and IUGR rats using the nitrate and nitrite colorimetric assay and the protein abundance of neuronal nitric oxide synthase (nNOS), endothelial NOS (eNOS), and phosphorylated endothelial NOS (peNOS) via Western blot analysis. To evaluate the amount of systemic and local amounts of NO, we measured the nitrate and nitrite, which are metabolites of NO, by using the R&D Systems Nitrate/Nitrite Assay Kit (780001, Cayman Chemical, Ann Arbor, MI) as described previously and according to the manufacturer’s instructions [36]. Optical Density (O.D.) measurements were read at 540 nm and corrected at 690 nm, to calculate total NOx in plasma (μM) and organ (μM/ug protein). All organ NOx amounts were normalized to total protein using a BCA assay.

### Oxidative Stress Profile Measurements

To evaluate systemic and localized levels of OS, we examined the plasma, kidney, and brain samples of pregnant CON and IUGR rats, measuring 8-isoprostanes, hydrogen peroxide (H_2_O_2_), and antioxidant capacity levels via biochemical assays as described by us previously [37]. See **Supplemental Section** for further details on methods.

### Western Blot Analysis for NO Bioavailability and Oxidative Stress

Western blots were used to identify and quantify specific proteins in the kidneys and brain pertaining to NO bioavailability (peNOS, nNOS, and eNOS) and OS (HSP-1, Cu/ZnSOD, MnSOD, ecSOD). For more details, please see the **Supplemental Methods Section**.

## STATISTICAL ANALYSIS

All data measurements are expressed as mean ± SEM. For comparing BP in groups (CON female x CON male, CON female x IUGR male, IUGR female x CON male, and IUGR female x IUGR male) using a two-way ANOVA with Bonferroni post hoc test. For all other comparisons of IUGR vs CON Preg, we used the unpaired two-tailed student t-test. Statistical significance was defined as a value of *p<0.05.

## RESULTS

### Maternal Blood Pressure, Body Weight, Organ Weight, and Fetal Weight

MAP was elevated in pregnant IUGR vs. CON rats (116.0 ± 4.2 vs. 100.6 ± 2.5 mmHg, p<0.05) (**Table 1**; **Figure 1A-B**). The MAP of the pregnant IUGR and CON rats was not affected by the status of the male rats (IUGR vs. CON), that were mated with the female IUGR and CON rats (**Figure 1B**). In other words, IUGR or CON males that were mated with female rats, had no influence on maternal BP. Furthermore, there was no difference in BP between IUGR and CON rats in the absence of pregnancy (**Supplemental 2**). This indicates that the increase in BP from pregnant IUGR rats was not due to the elevation of BP before pregnancy, but rather occur during pregnancy. There was no change in gestational day 19 body weight, heart weight, kidney weight, and placental weight (**Table 1**). However, there was a significant 9% increase in total brain weight in pregnant IUGR rats (5.4 ± 0.1 vs. 5.0 ± 0.2g/1000g BW, p<0.05) (**Table 1**). Nevertheless, a ∼20% reduction was observed with the average pup weight in pregnant IUGR rats (1.7 ± 0.1 vs. 2.1 ± 0.3g/1000g BW, p = 0.051) (**Table 1**).

**Figure 1.**
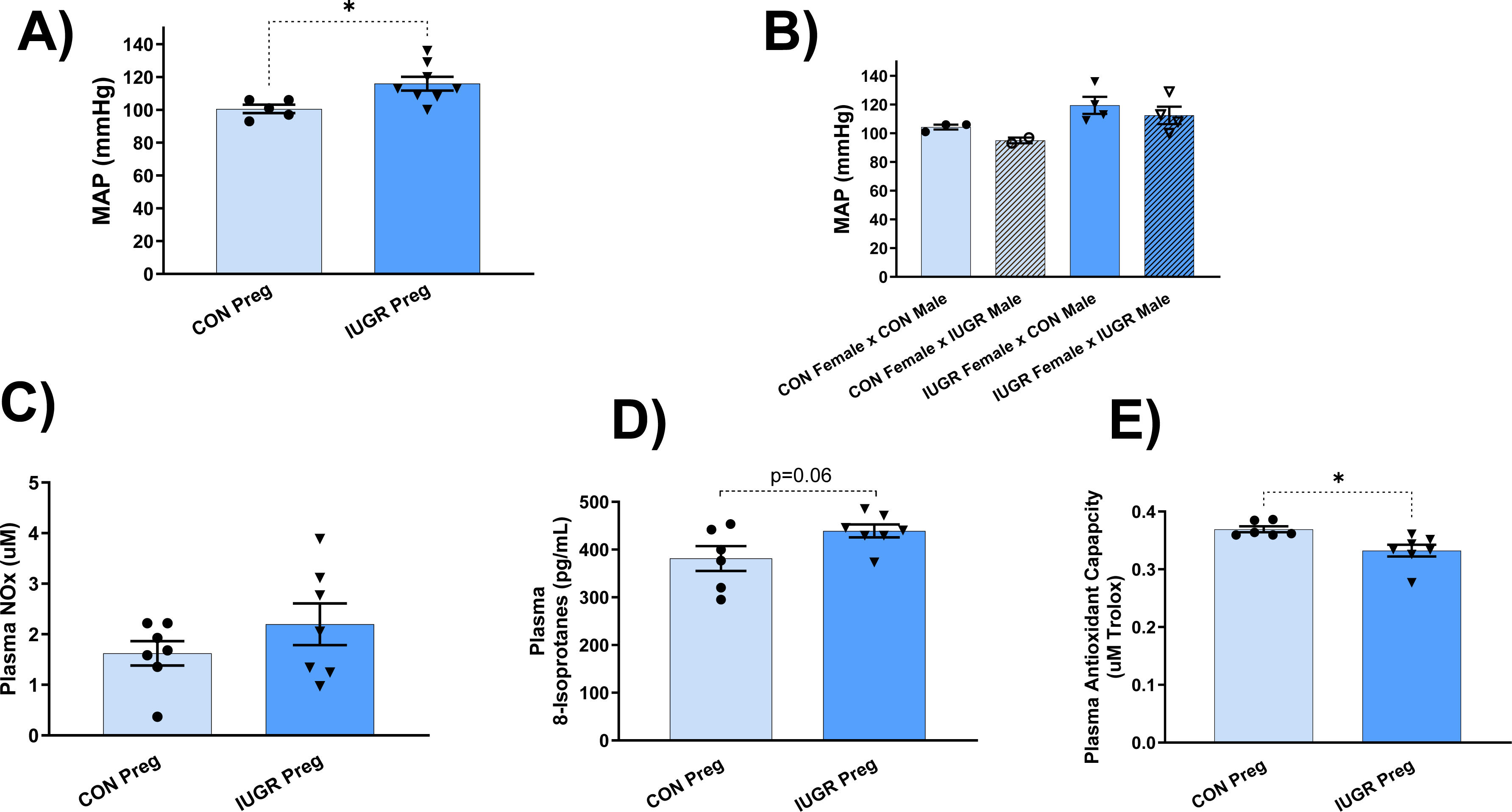
A) Examines mean arterial pressure (MAP) between pregnant IUGR (n=8) and CON (n=5) rats. **B)** Compares BP of female CON and IUGR rats mated with male CON and IUGR rats. Therefore, this figure is composed of 4 groups, which are: CON female x CON male, CON female x IUGR male, IUGR female x CON male, IUGR female x IUGR male. **C-E)** Compares plasma NOx, 8-isoprostanes (marker of systemic ROS), and antioxidant capacity respectively in pregnant IUGR (n=7) vs CON (n=6-7) rats. Significance, with a p-value < 0.05 vs CON, is indicated by an asterisk *. Statistical significance was determined in Figure 1B by two-way ANOVA, Bonferroni post hoc test and by student’s t-test in the other figures (Figure 1A**, C-E**) by comparing pregnant IUGR to CON rats.

**Table 1:** Displays maternal changes in BP, body weight (grams), heart weight (g/1000g body weight), kidney weight (g/1000g body weight), brain weight (g/1000g body weight), placental weight (g body weight), and average pup weight at gestational day 19 of pregnant IUGR (n=8) and CON (n=5) rats. Significance, with a p-value < 0.05 vs CON, is indicated by an asterisk *. Statistical significance was determined by student’s t-test comparing pregnant IUGR to CON rats.

|  | **CON Preg**<br>**(n= 5-6)** | **IUGR Preg**<br>**(n=8-9)** |
| --- | --- | --- |
| **Blood Pressure (mmHg)** | **100.6 ± 2.54** | **116.0 ± 4.17\*** |
| **GD 19 Body Weight (g)** | **350.3 ± 10.82** | **330.1 ± 5.24** |
| **Heart Weight (g/1000g BW)** | **2.58 ± 0.12** | **2.60 ± 0.05** |
| **Kidney Weight (g/1000g BW)** | **5.05 ± 0.18** | **5.07 ± 0.07** |
| **Brain Weight (g/1000g BW)** | **4.95 ± 0.19** | **5.38 ± 0.10\*** |
| **Placental Weight (g)** | **5.87 ± 0.84** | **6.27 ± 0.44** |
| **Average Pup Weight (g)** | **2.08 ± 0.27** | **1.69 ± 0.14** |

### Systemic and Local Evaluation of NO Bioavailability and Oxidative Stress. Plasma

There were no changes in plasma NOx levels in pregnant IUGR rats (2.2 ± 0.4 vs. 1.6 ± 0.2 μM, NS) (**Figure 1C**). Plasma 8-isoprostane (marker for ROS) levels trended upward in pregnant IUGR rats (381.3 ± 26.1 vs. 439.2 ±13.6 pg/mL, p=0.06) (**Figure 1D**). Plasma antioxidant capacity was lower in pregnant IUGR rats (0.3 ± 0.0 vs. 0.4 ± 0.0 μM Trolox, p<0.05) (**Figure 1E**).

### Kidney

To measure kidney NO bioavailability, we examine the protein levels of endothelial NOS (eNOS) and phosphorylated endothelial NOS (peNOS), the enzymes that generate NO production, along with NOx (nitrate and nitrite, metabolites of NO) production. No changes were observed in eNOS and peNOS protein abundance and NOx levels with pregnant IUGR rats **(Figure 2A-C)**. OS was evaluated in the kidney cortex by examining H_2_O_2_ (marker of ROS) levels, copper/zinc superoxide dismutase (Cu/ZnSOD; cytosolic SOD, antioxidant), manganese superoxide dismutase (MnSOD; mitochondrial antioxidant), and antioxidant capacity. No significant changes were observed in H_2_O_2_ levels, Cu/ZnSOD and MnSOD protein abundances, and antioxidant capacity **(Figure 3A-D)**.

**Figure 2.**
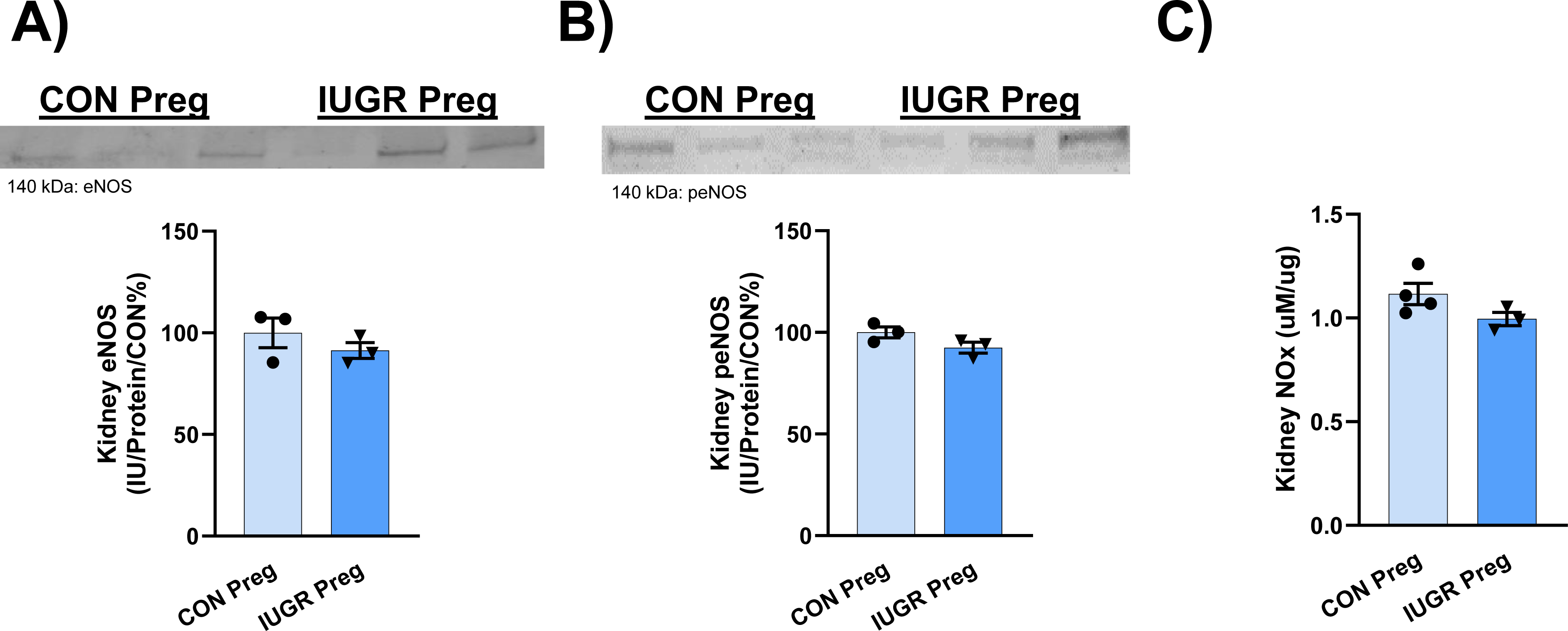
Kidney cortex NO bioavailability was evaluated by examining **A**) eNOS and **B**) peNOS protein abundance, via a Western blot image and quantitative bar graph of the image densitometry of pregnant IUGR (n=3) and CON (n=3) rats; along with **C**) NOx concentrations in pregnant IUGR (n=3) vs CON (n=4) rats. Significance, with a p-value < 0.05 vs CON, is indicated by an asterisk *. Statistical significance was determined by student’s t-test comparing pregnant IUGR to CON rats.

**Figure 3.**
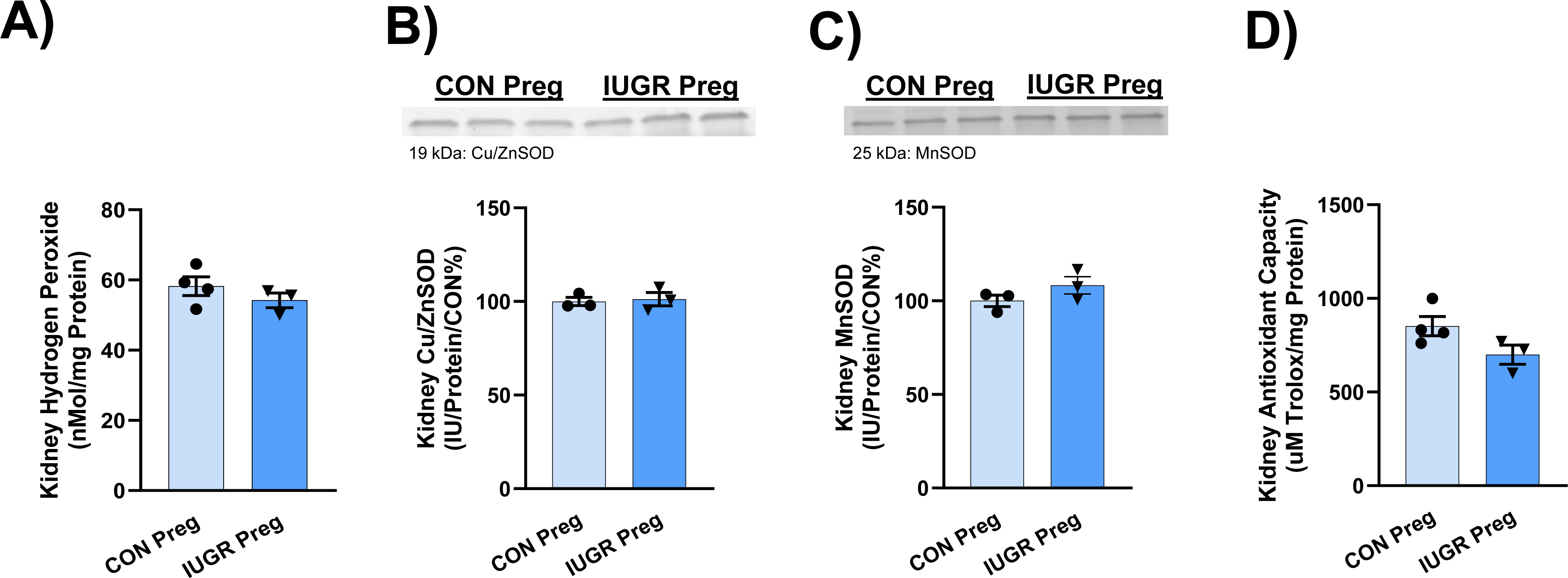
A) H_2_O_2_, as a marker of ROS, was measured in the kidney cortex of pregnant IUGR (n=3) and CON (n=4) rats. **B-C**) Antioxidant protein abundance of kidney cortex Cu/ZnSOD and MnSOD, is respectively displayed as a blot image and quantitative bar graph of the image using densitometry of pregnant IUGR (n=3) and CON (n=3) rats. **D**) kidney cortex antioxidant capacity of IUGR (n=3) and CON (n=4) pregnant rats. Significance, with a p-value < 0.05 vs CON, is indicated by an asterisk *. Statistical significance was determined by student’s t-test comparing pregnant IUGR to CON rats.

### Brain

No changes in brain neuronal NOS (nNOS) protein abundance and NOx were observed in pregnant IUGR rats (**Figure 4A-B**). However, H_2_O_2_ (marker of ROS) concentration was drastically increased, ∼ 2.4-fold, in pregnant IUGR rats (25.8 ± 5.1 vs. 11.4 ± 3.0 nMol/mg protein, p<0.05) (**Figure 5A**). Furthermore, heat shock protein 1 (HSP-1; marker of ROS) was substantially greater in protein abundance by ∼86% in pregnant IUGR rats (186 ± 28 vs. 100 ± 6 IU/Protein/CON %, p<0.05) (**Figure 5B**). There was no change in brain Cu/ZnSOD (**Figure 5C**), but there was a downward trend in brain MnSOD in pregnant IUGR rats (88 ± 3 vs 100 ± 3 IU/Protein/CON%, p=0.0540) (**Figure 5D**). There was no difference in brain antioxidant capacity in pregnant IUGR rats (260± 33 vs. 292± 14 μM Trolox/mg protein, NS) (**Figure 5E**).

**Figure 4.**
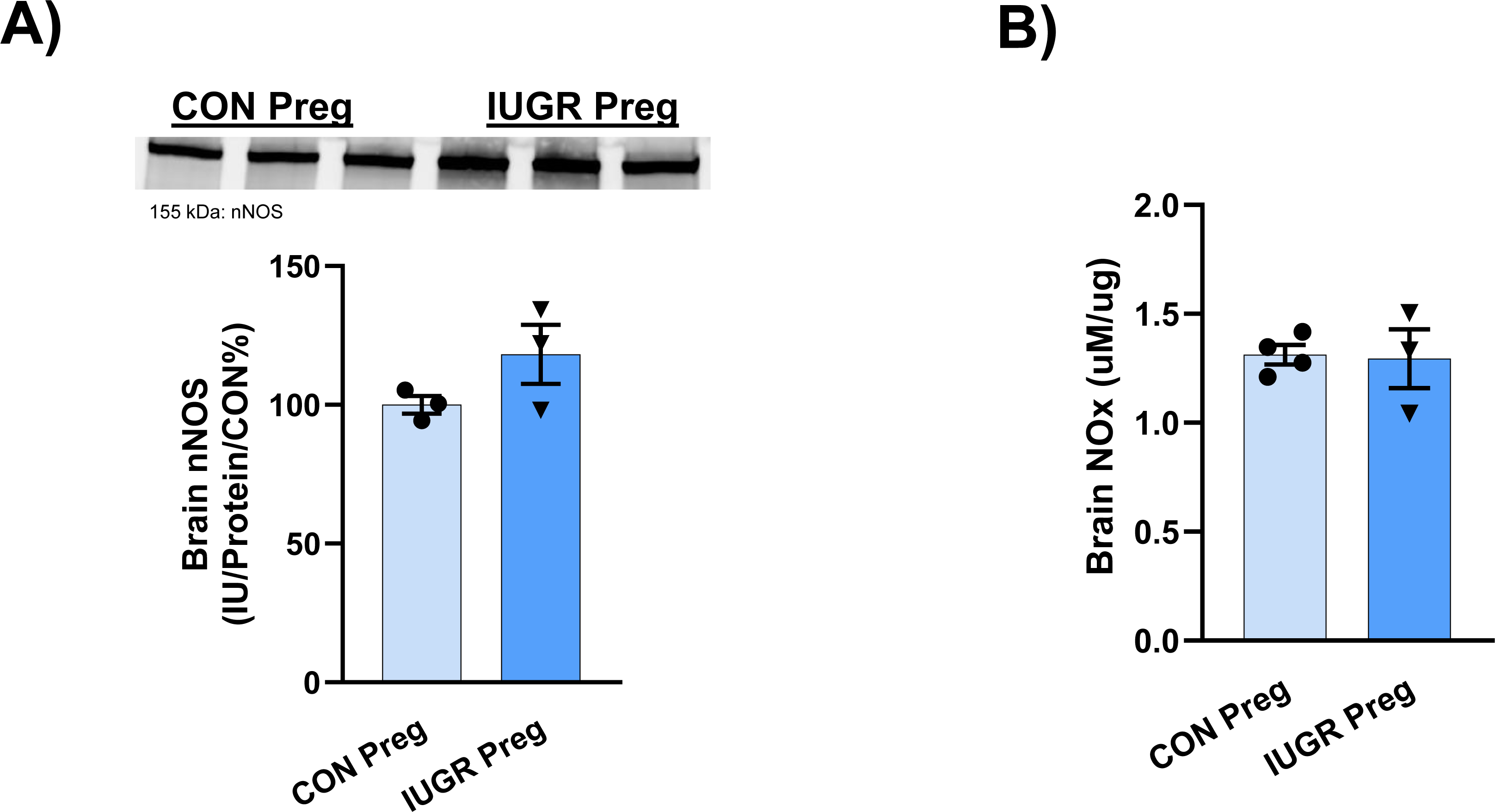
Brain NO bioavailability was elevated by examining **A**) nNOS, via a Western blot image and quantitative bar graph of the image below using densitometry of pregnant IUGR (n=3) and CON (n=3) rats and **B**) NOx (nitrate and nitrite) levels in pregnant IUGR (n=3) vs CON (n=3) rats. Significance, with a p-value < 0.05 vs CON, is indicated by an asterisk *. Statistical significance was determined by student’s t-test comparing pregnant IUGR to CON rats.

**Figure 5.**
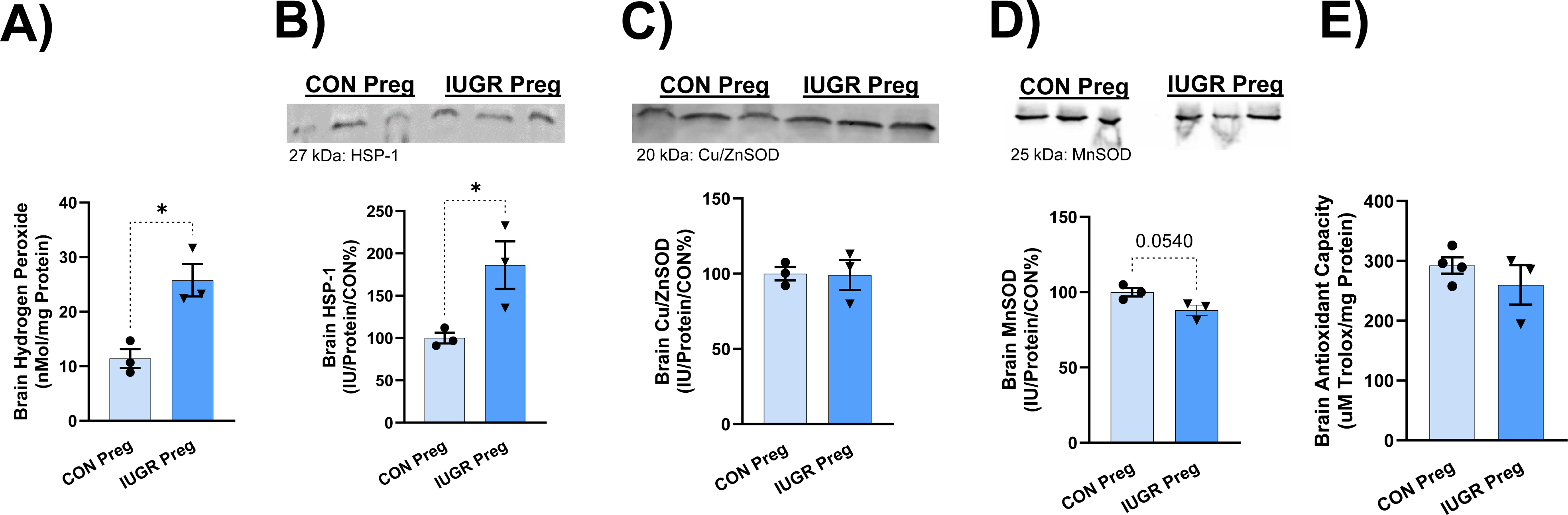
A) H_2_O_2_, as a marker of ROS, were measured in the brain of pregnant IUGR (n=3) and CON (n=3) rats. **B)** Brain HSP-1 (marker of ROS) is displayed as a representative blot image and quantitative bar graph of the image using densitometry of pregnant IUGR (n=3) and CON (n=3) rats. **C-D**) Antioxidant protein abundance of brain Cu/ZnSOD and MnSOD, is displayed, respectively, as a blot image and quantitative bar graph of the image using densitometry of pregnant IUGR (n=3) and CON (n=3) rats. **D**) Brain antioxidant capacity of pregnant IUGR (n=3) and CON (n=4) rats. Significance, with a p-value < 0.05 vs CON, is indicated by an asterisk *. Statistical significance was determined by student’s t-test comparing pregnant IUGR to CON rats.

## DISCUSSION

Pregnant IUGR rats, born from hypertensive placental ischemic (RUPP) dams with PE, have HTN and smaller pups at late gestation (day 19). In addition, they have elevated systemic and cerebral OS as well as larger brain sizes [38], which may lead to cerebral damage and/or injury commonly observed in preeclamptic women. Despite the cerebral damage that may be associated with this model, the kidneys appear to be less damaged or protected, showing no differences in NO bioavailability and OS. Therefore, more observations are needed to determine if there are dysfunctional changes in the renal vasculature, structure, and function. Whether or not HTN in these pregnant IUGR rats is due to cerebral damage and/or the cerebral OS in this model, is not yet known and will be the focus of future studies. However, pregnant IUGR rats from hypertensive placental ischemic rat dams, in this study, show symptoms of a preeclamptic- like phenotype, such as increased BP, low birth weight offspring, and increased OS. Thus, suggesting a new model to study PE.

PE is hypothesized to be caused by abnormal trophoblast invasion in the mother’s endometrium during pregnancy [11]. The decrease in trophoblast invasion causes abnormal placental spiral artery remodeling during placentation that results in a deficit of oxygen and nutrient exchange between the mother and the fetus. The net result of these abnormalities causes placental ischemia, endothelial dysfunction, and OS [4].

Both the mom and fetus have short and long-term health risks associated with the PE [1, 4, 39]. Furthermore, daughters of women with PE, have increased risk of having PE themselves [6]. In 2001, a genetic study performed in Utah found that PE is influenced by both maternal and paternal genes. However, the maternal influence was greater on the pathological development of PE [40]. Our studies mirror these findings and illustrated that birthing status (IUGR or CON) of the pregnant dam (mother) was a greater predictor of HTN during pregnancy (the major requirement for PE), regardless of the paternal birthing status (IUGR vs CON). Skjaerven et al., revealed that daughters born of a preeclamptic pregnancy exhibit a twofold greater risk of developing PE themselves [5]. In a more recent study, Sherf et al. concluded if the mom was an IUGR baby or had PE, then daughters had a higher incidence of PE in their pregnancies [6]. In summary, our data along with data in the literature from rodents and human studies, show that pregnant IUGR offspring display preeclamptic-like symptoms during pregnancy.

More specifically, few studies have examined pregnant rodents from second generation preeclamptic pregnancies [22–24]. Studies by Gallo et al. have shown that aged 12-month Wistar Kyoto restricted (IUGR) female rats from placental ischemic dams have elevated BP, before and during pregnancy, with normal renal and metabolic pregnancy adaptations. However, these aged, restricted rats display a normal drop in BP during pregnancy [23]. In addition, when they observed younger 4-month non-pregnant and pregnant restricted rats, they saw no differences in BP between non-pregnant and pregnant restricted and CON rats. Similarly, our IUGR 2.5-month Sprague Dawley rats from placental ischemic dams have no difference in BP before pregnancy in comparison to non-pregnant CON rats (See **Supplemental Figure 2**). Furthermore, our pregnant IUGR rats have an elevated BP of ∼ 16 mmHg higher than pregnant CON rats (**Figure 1A**). In addition, our pregnant IUGR rats in this study do not exhibit the normal drop in BP. However, the pregnant CON rats show the expected decrease in BP during pregnancy.

Although there are several other models of PE, they are all derived from either genetic and selective breeding (BPH mice and Dahl salt sensitive rat models), pharmacological (sFlt-1, AT-1-AA, L-NAME, and LPS models), and/or surgical interventions (RUPP model) [11, 16, 18–21]. Note that the soluble FMS-like tyrosine kinase 1 (sFlt-1) model, angiotensin 2 type 1 receptor autoantibody (AT1-AA) model, RUPP, L-Nitro-arginine methyl ester (L-NAME), and low-dose lipopolysaccharide (LPS) models are all animal models that express HTN, produce IUGR offspring, have decreased NO bioavailability, mitochondrial dysfunction, and elevated levels of OS, inflammation, and anti-angiogenic factors [39]. Whereas the inflammatory models of PE, such as tumor necrosis factor α (TNF-α) and interleukin 17 (IL-17) contain all previous characteristics, but varied results in producing IUGR offspring [11, 18]. The BPH mouse and Dahl salt sensitive (SS) rat models produce HTN, IUGR offspring, and elevated levels of OS, inflammation, and anti-angiogenic factors. Although the BPH and Dahl SS rodent models have provided a multitude of information on the pathology of PE, these models still require intervention via selective breeding. Both the BPH mouse and Dahl salt sensitive (SS) rat models were derived from selective breeding protocols to select the specific subtypes of hypertension in rodents. These hypertensive rodents were then selected and mated to generate the desired hypertensive phenotype of the rodents before and/or during pregnancy. Moreover, these models also produce rodents with metabolic dysfunction, obesity, increased adiposity, and/or borderline HTN before pregnancy [41]. In the Dahl SS rat model, when a high-salt diet is administered, the non-pregnant Dahl SS rat experiences an increase in BP, that continues to rise, during pregnancy. Thus, one may conclude that the HTN and preeclamptic-like phenotype in these rodent models are a continuation of an increase in BP from the non-pregnancy state, i.e. models of chronic HTN with superimposed PE. However, this is not the case with our rats, as shown by IUGR rat offspring that have no differences in BP without pregnancy (**Supplemental Figure 2**). Thus, our IUGR female rats are normotensive, expressing no difference in BP in comparison to CON rats without pregnancy. This observation has been investigated in other studies, such as by Alexander et al, that show IUGR female rats derived from the placental ischemic RUPP model of PE, after puberty (after 8 weeks of age), do not have elevated BP compared to female CON rats [11, 42]. However, the male IUGR rats have increased BP, and these sex differences may be due to the role of estrogen [43].

It is important to note that our non-pregnant female IUGR rats at 11-12 weeks of age did not have an increase in BP, changes in circulating glucose levels, nor changes body weight compared to non-pregnant female CON rats (**Supplemental Figure 2 and 3**). Note that we did not measure our pup weight of these IUGR offspring below 4 weeks of weaning for the fear of maternal rejection and rough handling of the offspring. Conversely, after 3 weeks of weaning, the offspring were closely observed for changes in weight, metabolic measurements (such as glucose), and BP until 11-12 weeks of age (**Supplemental Figure 1 and 2**). Thus, based on our data, it is reasonably assumed that our female IUGR rats did not have chronic HTN before pregnancy, but rather developed preeclamptic-like characteristics, such as HTN and the birth of IUGR offspring, during pregnancy.

Renal damage, as seen by proteinuria, elevated OS, and low NO bioavailability are common markers in preeclamptic animal models and humans [19, 44–46]. However, PE is a syndrome with a variety of possible symptoms and organ dysfunctions. For a pregnancy to be classified as preeclamptic, renal damage is not required. As of 2013, proteinuria, although a helpful diagnostic tool, does not alone reflect the severity or presence of PE in accordance with the American College of Obstetricians and Gynecologists [3, 47–49]. In our study, HTN and cerebral OS were present in pregnant IUGR rats, but the kidneys appear to be less damaged or protected, as observed by no changes in NO bioavailability and renal OS. Perhaps this protection is due to an increase and/or no loss in antioxidant capacity at this age, but more tests are needed to affirm this response. The kidneys, as a major regulator of BP, in the pregnant IUGR rat do not seem to play a prominent role in the pathophysiology of HTN and developments of preeclamptic-like symptoms in our model. Other markers of renal damage such as decreased glomerular filtration rate and increased albumin, proteinuria, creatinine, and vascular tone were not tested in this study, but are needed for renal analyses.

An important contributor to BP regulation is the brain. The common occurrence of cerebral symptoms reveals that preeclamptic women are vulnerable to brain damage and abnormalities [1, 50–52]. The increase in the BP and preeclamptic-like pathophysiology could be caused or facilitated by cerebral damage. The link between cerebral damage and HTN is unknown in this model and will be addressed in future studies. To determine the mechanisms and development of HTN and cerebral damage, we will conduct timeline and drug manipulation experiments [49]. However, our data supports current studies indicating the presence of increased cerebral OS and edema in PE patients [52–54]. Since the brain is implicated in the pathophysiology of PE, perhaps brain-directed therapies to increase antioxidant capacity and reduce ROS could ameliorate the PE symptoms.

Although this study examines some of the markers associated with PE, there are many more to evaluate in the future. For example, mitochondrial dysfunction, pro-inflammatory factors/cells (Il-6, IL-17), anti-angiogenic factors (sFlt-1, soluble endoglin), and placental ischemia [7, 32]. Placental ischemia is one of the major hallmarks of PE, and unfortunately, we did not have the proper equipment to make these measurements in this study. Furthermore, vasculature resistance needs to be evaluated to determine its possible role in the increase in BP displayed in our pregnant IUGR offspring. Studies have shown that female IUGR rodent offspring in the absence of pregnancy have vascular dysfunction [55, 56]. However, whether these IUGR offspring with vascular dysfunction have elevated BP or not is controversial. Some studies show no change in BP in IUGR offspring with only uterine vascular dysfunction [56]; while others show mesenteric vascular dysfunction, with a special emphasis in vasoconstrictor responses, and BP elevation in both first and second-generation IUGR offspring [55, 56].

## CONCLUSION

In conclusion, this study demonstrates that female IUGR rats from hypertensive PE ischemic dams have preeclamptic-like symptoms during pregnancy. These pregnant IUGR rats have HTN, elevated systemic and cerebral OS, enlarged brains, and smaller pups during late gestation, which are all clinical observations of women with PE. Unlike previous PE animal models that utilize selective breeding protocols to induce PE, our model uses pregnant IUGR rats as another viable method to study PE, especially regarding its effects on pregnant daughters from PE mothers. Changes in cerebral OS observed in pregnant IUGR rats suggest that antioxidant- specific and/or organ-targeted therapy may be beneficial to PE women, although more studies are needed.

Findings from this study are important to shift focus towards preventative care, especially in at-risk populations of PE, in which the mother had PE. Thus, the importance of pregnant women knowing their mother’s pregnancy complications and/or their IUGR status at birth maybe important for the development of PE during their pregnancy. This study also advises physicians to carefully examine populations of women that were born IUGR and whose mother was preeclamptic before and during their pregnancy. Based off this study, we advocate that all pregnant women should know their mother’s pregnancy history, because it may be beneficial to their overall health.

## PERSPECIVE/CLINICAL SIGNIFICANCE

This study is clinically significant because it supports the notion that complicated hypertensive pregnancies, such as preeclampsia, have long-lasting effects on the offspring, especially in pregnant daughters of women with PE. Furthermore, pregnant women need to know their birthing conditions and PE status of their mother, because their mother’s birthing condition could affect their own pregnancy. Information shared by the mother could be used by physicians and scientists to give them insights into the pathophysiology, risk, and prevention of PE in women that are IUGR and/or mother’s had PE.

## ACKNOWLEDGMENTS

The authors acknowledge the support of the Department of Physiology and Anatomy at the University of North Texas Health Science Center.

## ABBREVIATIONS

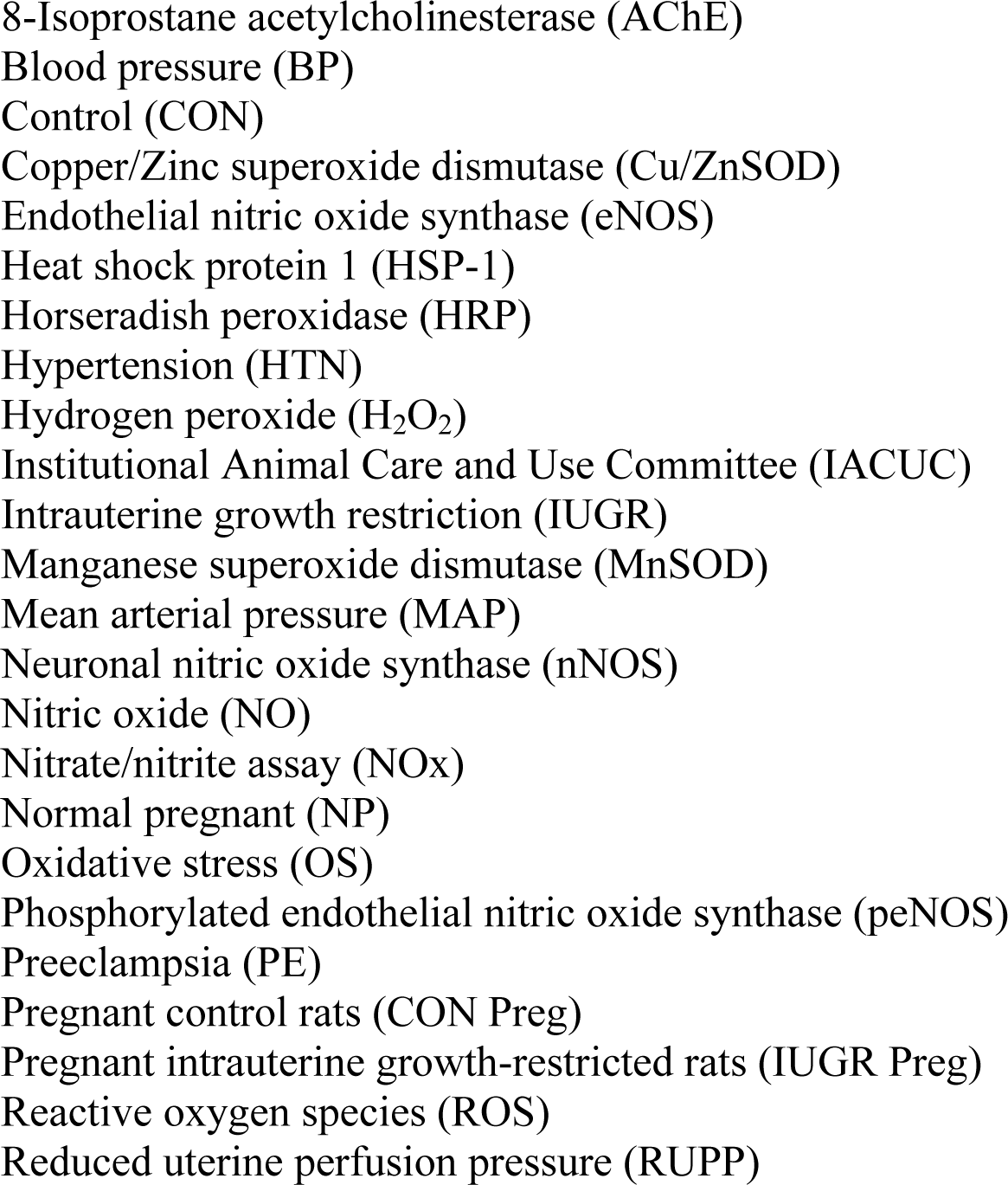

